# Dental caries in wild primates: interproximal cavities on anterior teeth

**DOI:** 10.1101/2021.06.30.450645

**Authors:** Ian Towle, Joel D. Irish, Kristin Sabbi, Carolina Loch

**Author notes:** **Corresponding author contact**: Ian Towle, Postal address: Walsh Building, North Dunedin, Dunedin 9016.

## Abstract

Dental caries has been reported in a variety of primates, although is still considered rare in wild populations. In this study, 11 catarrhine primates were studied for the presence of caries. A differential diagnosis of lesions found in interproximal regions of anterior teeth was undertaken, since they had been previously described as both carious and non-carious in origin. Each permanent tooth was examined macroscopically, with severity and position of lesions recorded. Two specimens were micro-CT scanned to assess demineralization. The differential diagnosis confirmed the cariogenic nature of interproximal cavities on anterior teeth (ICAT’s). Overall results show 3.3% of teeth are carious, with prevalence varying among species from 0% to over 7% of teeth affected. ICAT’s occurred in Pan troglodytes (9.8%), *Gorilla gorilla gorilla* (2.6%), *Cercopithecus denti* (22.4%), *Presbytis femoralis* (19.5%) and *Cercopithecus mitis* (18.3%). They make up 87.9% of carious lesions on anterior teeth. These results likely reflect dietary and food processing differences among species, but also between the sexes (e.g., 9.3% of teeth of female chimpanzees were carious vs. 1.8% in males). Processing cariogenic fruits and seeds with the anterior dentition (e.g., wadging) likely contributes to ICAT formation. Further research is needed on living populations to ascertain behavioral/dietary influences on caries occurrence in primates. Given the constancy of ICAT’s in frugivorous primates, their presence in archaeological and paleontological specimens may shed light on diet and food processing behaviors in fossil primates.

## Introduction

Caries formation is influenced by dietary, behavioral, environmental, and genetic factors (Kotecha et al., 2012; Slade et al., 2013). However, such lesions ultimately form from acids produced by cariogenic bacteria metabolizing sugars and starches, leading to demineralization of dental hard tissues (Byun et al., 2004; Larsen et al., 1991). Several microorganisms have been implicated in this process, including *Streptococcus sobrinus* and *S. mutans* (Nishikawara et al., 2007). The composition of the oral biofilm is a key component in caries formation (Cornejo et al., 2013); however, the same bacteria are often a normal part of the oral microbiome (Aas et al., 2008; Simón-Soro and Mira, 2015). Therefore, most primates likely have the potential for caries formation if enough cariogenic foods are consumed (Sheiham and James, 2015).

Caries research has historically focused on humans, with high prevalence of lesions often associated with an agriculturalist lifestyle, or hunter-gatherer populations that consume specific cariogenic foods (Caglar et al., 2007; Lanfranco & Eggers, 2012; Esclassan et al., 2009; Novak, 2015; Slaus et al., 2011; Srejic, 2001; Varrela, 1991; Watt et al., 1997; Walker and Hewlett, 1990; Sealy et al., 1992; Humphrey et al., 2014; Nelson et al., 1999). In non-agricultural hominins, typically less than 5% of teeth are carious (Towle et al., 2021; Turner, 1979; Lacy, 2014; Kelley et al., 1991; Larsen et al., 1991). In these groups, foods containing high levels of carbohydrates are implicated in caries formation, whereas tough and fibrous foods are often linked with low caries rates because of high wear rates and increased saliva production (Clarkson et al., 1987; Moynihan, 2000; Novak, 2015; Prowse et al., 2008; Rohnbogner & Lewis, 2016). Diets rich in fruits, seeds, and nuts are often associated with high rates, with varying susceptibility depending on the type of foods and how they are processed orally (Humphrey et al., 2014; Novak, 2015). A variety of other factors can affect the likelihood of caries, including the extent and type of crown wear and other pathologies/defects, such as enamel hypoplasia and periodontal disease (Hillson, 2008; Calcagno and Gibson, 1991; Towle and Irish, 2019; Towle and Irish, 2020).

Non-human primates also develop caries, particularly in captivity, and lesions have been described in extant and extinct wild populations (e.g., Cohen and Goldman, 1960; Coyler, 1936; Fuss et al., 2018; Lovell, 1990; Miles & Grigson, 2003; Schultz, 1935; 1956; Smith et al., 1977; Stoner, 1995). In humans, posterior teeth are most commonly affected. In non-human primates, anterior teeth evidence higher rates (Colyer, 1931), though in captivity the pattern changes to posterior teeth (Anderson & Arnim, 1937; Bowen, 1968; Cohen & Bowen, 1966; Colyer, 1936). Typically, lesions on anterior teeth form in interproximal regions of incisors (Schultz, 1935; Smith et al., 1977; Stoner, 1995; Lovell, 1990). These Interproximal cavities on anterior teeth (ICAT) have not always been regarded as carious, or otherwise have been overlooked (e.g., Kilgore, 1989).

Therefore, although ICAT’s have been previously reported, a study on their occurrence in a wide range of primate species is required. In this exploratory study, we use micro-CT scans, and consider other potential forms of tissue removal that may lead to ICAT formation. The influence of sex, age, and certain pathologies (e.g., abscesses and periodontal disease) on the formation of carious lesions was also considered. A total of 11 catarrhine species were selected, with an emphasis on frugivores, given the cariogenic potential of many fruits. We hypothesize that species known to regularly process sugary fruits with their anterior dentition, including behaviors such as ‘wadging’, will display ICATs, while those with a more varied diet will have a lower incidence.

## Methods

All samples studied here are curated at the Primate Research Institute, Kyoto University, Japan, and the Powell-Cotton Museum, UK. The 11 catarrhine species include: chimpanzee (*Pan troglodytes*), Western lowland gorilla (*Gorilla gorilla gorilla*), Kloss’s gibbon (*Hylobates klossii*), hamadryas baboon (*Papio hamadryas*), pig-tailed langur (*Simias concolor*), Japanese macaque (Macaca fuscata), Dent’s mona monkey (*Cercopithecus denti*), blue monkey (*Cercopithecus mitis*), mandrill (*Mandrillus* sp.), raffles’ banded langur (*Presbytis femoralis*), and Mentawai langur (*Presbytis potenziani*). All specimens were wild, and lived and died in their natural habitat (Buck et al., 2018; Guatelli-Steinberg and Skinner, 2000; Kamaluddin et al., 2019; Lukacs, 2001). The specimen numbers are presented in the supplementary material.

All permanent teeth retained within the jaws of each specimen were examined macroscopically. Those with substantial postmortem damage were excluded from analysis. Two teeth were subsequently removed for micro-CT scanning to ascertain enamel/dentine demineralization, and to visualize lesion progression (Boca et al., 2017; Rossi et al., 2004; Swain & Xue, 2009). Scans were performed at the Primate Research Institute, Kyoto University, using a SkyScan1275 Micro-CT scanner. The two teeth belonged to different Dent’s mona monkeys, one with a cavity on the mesial surface (upper right central incisor; PRI 11580) and the other displaying only coloration changes in the same location (PRI 11578; lower left central incisor).

X-rays were generated at 100 kV, 100 µA and 10W, with a 1mm copper filter placed in the beam path. Resolution was set at 15 µm pixel size, and rotation was set to 0.2-degree for both teeth. Images were reconstructed using the Skyscan NRecon software (NRecon, version 1.4.4, Skyscan) with standardized settings (smoothing: 3; ring artifact correction: 10; beam hardening: 30%). Resin-hydroxyapatite phantoms were used to calibrate greyscales and mineral densities in each specimen (Schwass et al. 2009). The calibration followed Schwass et al. (2009) and Loch et al. (2013). Data collection from the scans was undertaken using ImageJ. After calibration with phantoms, mineral concentration and total effective density was calculated for four locations in each tooth (buccal, lingual, distal and mesial). For the site with the potential carious lesion (mesial), three readings (oval ROI: 0.15mm diameter) were taken in 10 slices (total 30 measurements), with the slice interval based on the extent of the lesion. The individual ROI’s were chosen at random within a distance of 0.5mm of the interproximal surface of the dentine (i.e., directly adjacent to the cavity in PRI 11580 and beneath the area of coloration change in PRI 11578). The same data were collected for the other three locations (buccal, lingual and distal), on the same slices, with random ROI’s selected within 0.5mm of the dentine edge.

Caries prevalence by species was calculated as the percentage of carious teeth among all permanent teeth analyzed, including antimeres. Color changes on the dental tissues were not considered diagnostic, but were recorded when in association with antemortem cavitation. Cavity severity and position on the crown were also recorded. Severity was scored on a scale of 1 to 4 (Connell and Rauxloh, 2003): (1) enamel destruction; (2) compromised dentine but pulp chamber not exposed; (3) destruction of dentine with pulp exposure; (4) gross destruction with crown mostly destroyed. Location was recorded as buccal, occlusal, distal, lingual and mesial. If it was not possible to determine location due to severity, the lesion was recorded as ‘gross’. Due to difficulty in ascertaining if certain lesions would be best described as affecting the crown or root, teeth are not divided into these categories; however, potential differences are discussed.

Caries can directly contribute to antemortem tooth loss, which has led to correction methods (e.g., Duyar and Erdal, 2003; Kelley et al., 1991; Lukacs, 1995). The most likely causes of antemortem tooth loss in extant primates are severe attrition, fractures, and periodontal disease. Therefore, following other studies (e.g., Larsen et al., 1991; Meinl et al., 2010), no correction methods were implemented. Data on abscesses and periodontal disease were collected following Dias and Tayles (1997) and Ogden (2007), to assess if these pathologies were associated with caries. Each individual was recorded as having caries, periodontal disease, and abscesses (see Supplementary file). For abscesses and caries, the individual needed to show at least one lesion in the dentition to be recorded as affected. For periodontal disease, the individual needed to exhibit general resorption (rather than just pockets) in the mandible and/or maxilla.

Species were also divided by sex to explore differences in caries occurrence. Wear was scored following Scott (1979) for molars, and Smith (1984) for all other teeth. For sex difference analyses, a Chi-square test with alpha level of 0.05 was used.

## Results

Caries frequency was low (<1.5% of teeth) in over half the species. Posterior tooth caries was particularly rare, though some species showed a moderate prevalence (Table 1). Nearly 88% of all lesions on anterior teeth were interproximal, with mesial surfaces mostly affected (Table 2). These ICATs were only present in chimpanzees, Western lowland gorillas, Dent’s mona monkeys, blue monkeys, and raffles’ banded langurs. Most anterior lesions were small (severity 1), although the chimpanzee and gorilla samples showed higher frequencies of larger lesions (Table 3).

**Table 1.** Caries prevalence for permanent teeth for each species studied, split by anterior and posterior teeth. Figure in parenthesis is the percentage of ICAT teeth (i.e., anterior teeth with mesial and/or distal carious lesions).

| Species | Common name | Anterior teeth |  |  | Posterior teeth |  |  | All teeth |  |  |
| --- | --- | --- | --- | --- | --- | --- | --- | --- | --- | --- |
|  |  | # Teeth | carious teeth | % | # Teeth | carious teeth | % | # Teeth | carious teeth | % |
| *Pan troglodytes* | Chimpanzee | 914 | 112 | 12.3 (9.8) | 1584 | 53 | 3.4 | 2498 | 165 | 6.6 |
| *Gorilla gorilla gorilla* | Western lowland gorilla | 779 | 20 | 2.6 (2.6) | 1311 | 0 | 0 | 2090 | 20 | 1 |
| *Cercopithecus denti* | Dent's mona monkey | 85 | 19 | 22.4 (22.4) | 180 | 0 | 0 | 265 | 19 | 7.2 |
| *Cercopithecus mitis* | Blue monkey | 82 | 15 | 18.3 (18.3) | 160 | 1 | 0.6 | 242 | 16 | 6.6 |
| *Mandrillus sp.* | Mandrill | 20 | 0 | 0 | 108 | 2 | 1.9 | 128 | 2 | 1.6 |
| *Presbytis femoralis* | Raffles' banded langur | 82 | 16 | 19.5 (19.5) | 160 | 2 | 1.3 | 242 | 18 | 7.4 |
| *Presbytis potenziani* | Mentawai langur | 69 | 0 | 0 | 158 | 0 | 0 | 227 | 0 | 0 |
| *Hylobates klossii* | Kloss's gibbon | 84 | 0 | 0 | 232 | 4 | 1.7 | 316 | 4 | 1.3 |
| *Macaca fuscata* | Japanese macaque | 298 | 0 | 0 | 713 | 12 | 1.7 | 1011 | 12 | 1.2 |
| *Simias concolor* | Pig-tailed langur | 132 | 0 | 0 | 277 | 0 | 0 | 409 | 0 | 0 |
| *Papio hamadryas* | Hamadryas baboon | 182 | 0 | 0 | 336 | 6 | 1.8 | 518 | 6 | 1.2 |
| All species combined |  | 2727 | 182 | 6.7 | 5219 | 80 | 1.5 | 7946 | 262 | 3.3 |

**Table 2.** Percentage of carious lesions recorded for each surface on anterior teeth for the five species with anterior tooth lesions. Bold figures indicate position with highest caries prevalence.

| Species | Common name | Buccal | Mesial | Lingual | Distal | Gross | Occlusal |
| --- | --- | --- | --- | --- | --- | --- | --- |
| *Pan troglodytes* | Chimpanzee | 8.1 | **68.5** | 2.7 | 11.7 | 9 | 0 |
| *Gorilla gorilla gorilla* | Western lowland gorilla | 0 | **70** | 0 | 30 | 0 | 0 |
| *Cercopithecus denti* | Dent's mona monkey | 0 | **69.2** | 0 | 30.8 | 0 | 0 |
| *Cercopithecus mitis* | Blue monkey | 0 | **81.3** | 0 | 18.8 | 0 | 0 |
| *Presbytis femoralis* | Raffles' banded langur | 0 | **63.2** | 5.3 | 31.6 | 0 | 0 |

**Table 3.**
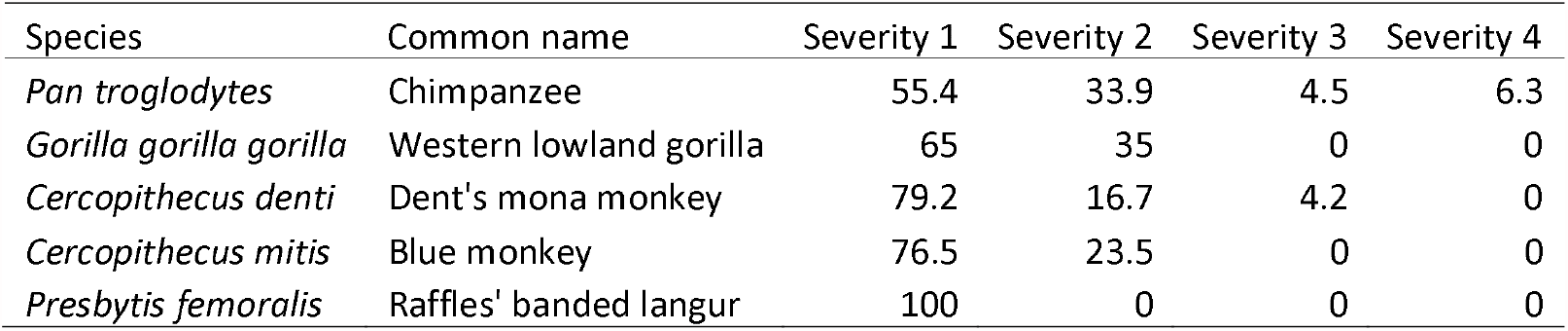
Percentage of carious lesions recorded for each severity grade on anterior teeth of species with interproximal lesions.

Micro-CT scans of two above mentioned teeth revealed that dentine was substantially demineralized beneath both the cavitation and color change areas (Figure 1). In both cases, a much lower mineral concentration and total effective density was evident compared to other areas of the tooth (Table 4). The range and standard deviation of the dentine mineral concentration within the carious locations was also much greater than in sound dentine, adding further support for demineralization caused by caries. Below the cavity in specimen PRI 11580, demineralization reached deep into the dentine, reaching approximately 0.5mm before tissue returned to normal density (i.e., over 1.6g/cm3).

**Table 4.** Average mineral concentration and total effective density (g/cm^3^) for each dentine position studied, along with standard deviation (±) and minimum and maximum values (in parenthesis), for a tooth with a large cavity (PRI 11580) and a tooth showing coloration changes but no cavitation (PRI 11578).

|  | Average Mineral Concentration | Average Total Effective Density |
| --- | --- | --- |
| **PRI 11580** |  |  |
| Mesial (lesion) | $0.60 \pm 0.12$ (0.43-0.85) | $1.43 \pm 0.05$ (1.36-1.52) |
| Buccal | $1.29 \pm 0.06$ (1.17-1.43) | $1.70 \pm 0.02$ (1.65-1.75) |
| Distal | $1.18 \pm 0.03$ (1.10-1.23) | $1.65 \pm 0.01$ (1.62-1.67) |
| Lingual | $1.17 \pm 0.04$ (1.10-1.24) | $1.65 \pm 0.02$ (1.62-1.68) |
| **PRI 11578** |  |  |
| Mesial (lesion) | $0.84 \pm 0.08$ (0.73-1.02) | $1.52 \pm 0.03$ (1.48-1.59) |
| Buccal | $1.20 \pm 0.03$ (1.13-1.26) | $1.66 \pm 0.01$ (1.63-1.69) |
| Distal | $1.20 \pm 0.01$ (1.17-1.23) | $1.66 \pm 0.01$ (1.65-1.68) |
| Lingual | $1.24 \pm 0.03$ (1.17-1.30) | $1.68 \pm 0.01$ (1.65-1.70) |

**Figure 1.**
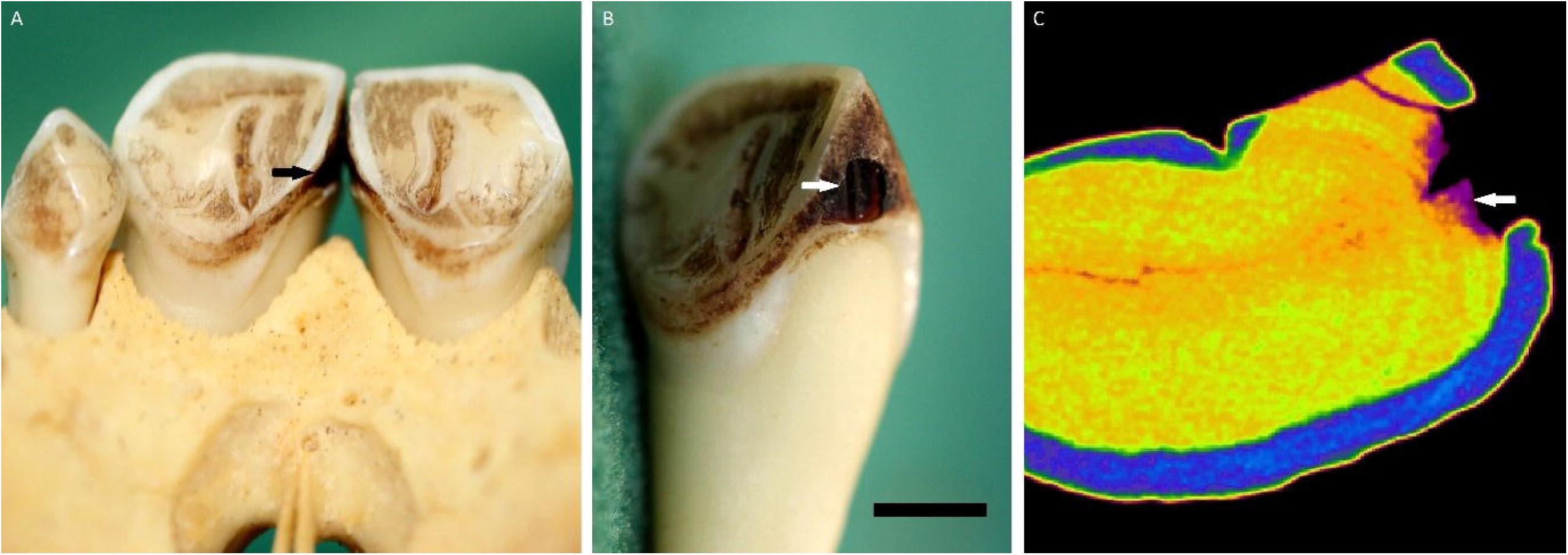
Carious lesions on the mesial surface of an upper right central incisor in a Dent’s mona monkey (*Cercopithecus denti*; PRI 11580) individual. A) carious lesion (black arrow) showing the relationship to the adjacent left central incisor; B) close up of the lesion (white arrow), scale bar 5mm; C) Micro-CT slice of the same lesion, showing demineralization deep into the dentine (white arrow).

The five species displayed ICATs that were similar in shape and crown position, with relatively circular lesions near the cementum-enamel junction (CEJ) on the interproximal surfaces of anterior teeth (Figures 2a). In more severe cases, much of the crown was destroyed (Figure 2b). In individuals with mild to moderate severity, lesions were limited to the CEJ region, and often only affected the crown; however, in some cases, lesions initiated on the root and followed the CEJ boundary, giving them a more oval appearance. In many cases it was not possible to ascertain if the lesion initially involved the crown or root, as the cavity covered both regions. Many individuals showed a dark coloration in these interproximal regions, but no cavitation (Figure 3). Along with the micro-CT scan data, this feature suggests these areas were early carious lesions.

**Figure 2.**
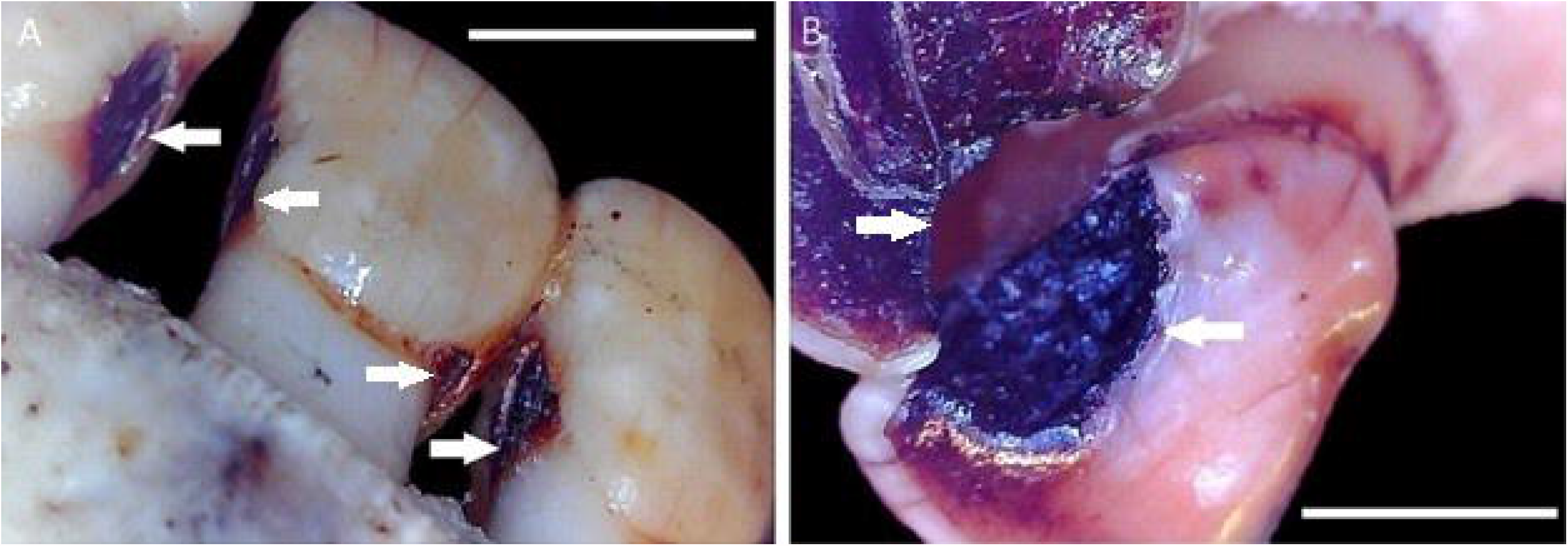
Interproximal caries in chimpanzees (*Pan troglodytes*): A) M60: carious lesions on mandibular left lateral incisor and both central incisors (indicated by white arrows); B) M155: carious lesions in maxillary right central and lateral incisors (indicated by white arrows). Both scale bars are 5mm.

**Figure 3.**
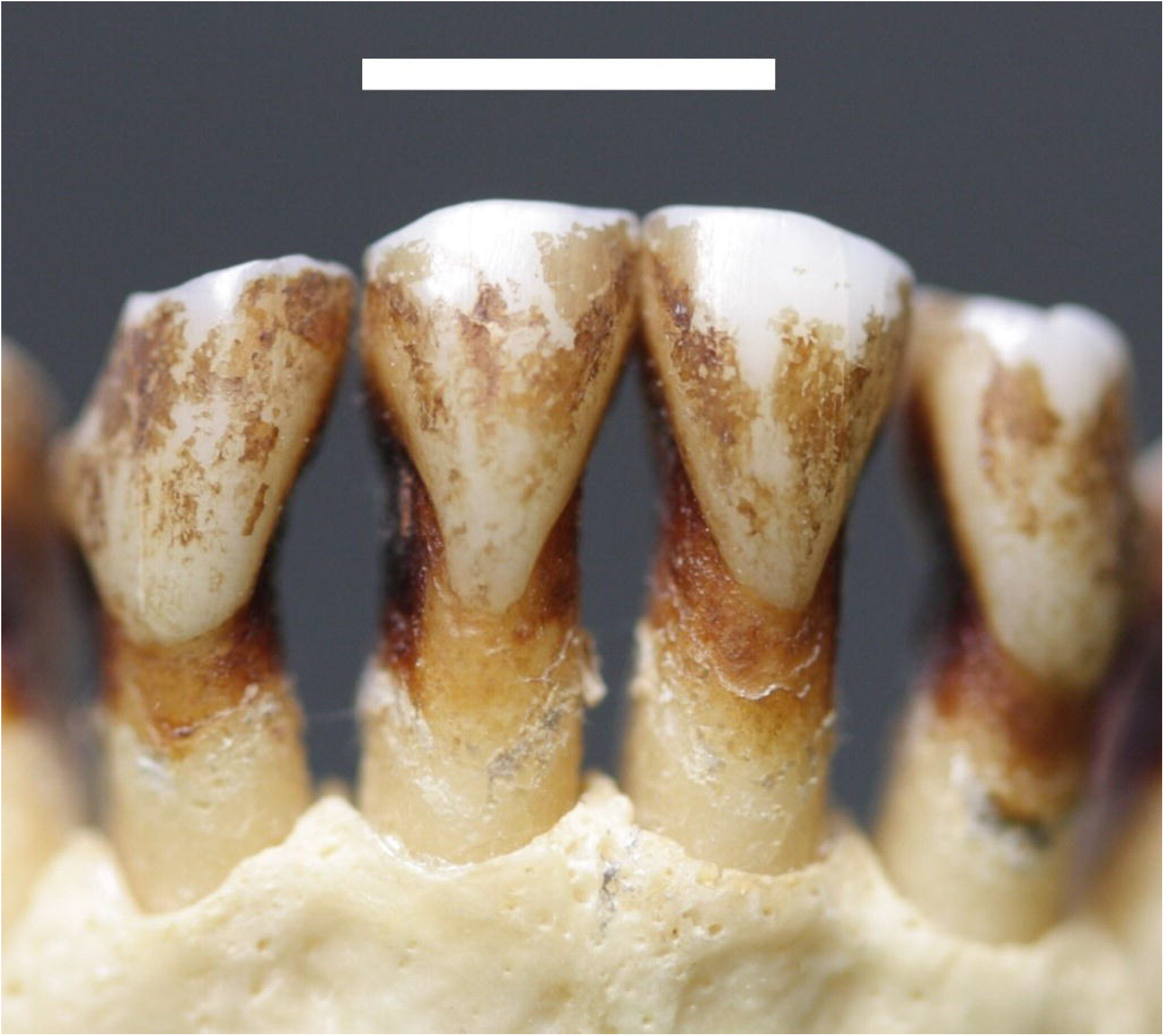
Raffles’ banded langur (*Presbytis femoralis*; PRI 4565) lower incisors displaying potential periodontal disease and early stages of caries in the interproximal areas, but no clear evidence of cavitation. Scale bar 5mm.

Periodontal disease and extensive occlusal/interproximal wear were sometimes associated with caries formation (Figs. 1 and 2). However, species with high caries levels do not seem to have an overall increase in periodontal disease, with both carious and non-carious individuals similarly affected. Small sample sizes for most species hampered statistical comparisons (Supplementary file).

Only chimpanzees displayed a significant difference in prevalence of caries between males and females (Supplementary Table 2). This analysis was hindered by small sample sizes for Cercopithecidae. However, when pooled together, there was little difference in caries frequency by sex (Cercopithecidae species combined; males: 25.49%; females: 17.57%; × = 1.1504, 1 df, p= 0.28). While a slightly higher percent of male gorillas have caries, this difference is not statistically significant (males: 11.43%; females: 6.52%; X^2^ = 0.6062, 1 df, p= 0.44). In contrast, female chimpanzees had significantly more caries than males (males: 17.07%; females: 40%; X^2^ = 6.1639, 1 df, p= 0.01).

When sex differences in caries occurrence were compared in terms of number of teeth affected, female chimpanzees also had more caries (X^2^ = 20.890, 1 df, p= 0.00), with five times the number of teeth affected. This difference does not appear to relate to age, based on crown wear (Table 5). Although most female chimpanzees exhibited more crown wear, this difference in caries frequency remained stable once teeth were split into wear categories. Females with low and medium levels of wear (combined wear score under 64 for all four first molars; following Scott, 1979) displayed more carious teeth, with five times the rate of males in the same wear category.

**Table 5.** Caries prevalence for male and female chimpanzees. Displayed for all teeth, unworn/little-worn teeth removed, and with heavily worn teeth excluded. I: incisors; C: canines; PM: premolars.

| **Sample** | **Females** | **Males** |
| --- | --- | --- |
| **Totals (%)** |  |  |
| Total teeth | 1301 | 334 |
| Carious teeth | 121 | 6 |
| Carious teeth % | 9.3 | 1.8 |
| % of individuals with caries | 44.9 | 8.3 |
| Mean I, C and PM wear** | 3.9 | 2.6 |
| Mean molar wear** | 4.1 | 2.7 |
| **Wear score 1 taken out** |  |  |
| Total teeth | 1192 | 255 |
| Carious teeth | 121 | 6 |
| Mean I, C and PM wear** | 4.31 | 3.5 |
| Mean molar wear** | 4.19 | 2.9 |
| % caries teeth | 10.2 | 2.4 |
| **Medium to low wear\*** |  |  |
| Total teeth | 511 | 227 |
| Carious teeth | 50 | 5 |
| Mean I, C and PM wear** | 3.5 | 3.6 |
| Mean molar wear** | 3.3 | 2.8 |
| % carious teeth | 9.8 | 2.2 |

\*Individuals with a combined wear score of under 64 for all four first molars. Teeth with a wear score of 1 are excluded
\*\*Molar wear is calculated using Scott (1979) and all other teeth following Smith (1984)

## Discussion

The results of this study suggest caries frequency was relatively low in the primates studied (0-7.4% of teeth; Table 1). Anterior teeth had a higher frequency, and lesions were similar among species in terms of position and physical characteristics. In particular, ICATs appear to be relatively common in frugivorous and seed eating Cercopithecidae and Hominoidea, likely related to the way dietary items are orally processed.

Although Kilgore (1989) suggested ICAT’s may relate to severe enamel attrition from stripping foods, demineralization visible deep into the dentine on the micro-CT scans is strongly suggestive of caries. Attrition-related behavior could contribute to lesion formation, by exposing the underlying dentine in interproximal areas. However, the present micro-CT scan results, radiographs in Kilgore (1989), and thin sections in Miles and Grigson (2003), seem to confirm that caries is the predominate factor for tissue loss in ICATs. Furthermore, coloration changes in these regions are suggestive of early stage demineralization associated with caries; thus, the true rate and effects of caries are likely much higher than the present findings suggest. Other processes, such as attrition or non-bacterial erosion, are unlikely to yield localized deep cavities and coloration change visible in ICATs.

Other researchers have reported ICATs as carious lesions (e.g., Colyer, 1936; Schultz, 1956). These studies also observed high caries rates in chimpanzees, with incisors commonly affected. Colyer (1931) reported that in wild monkeys, anterior teeth were more commonly affected than posterior teeth (66.2% vs. 33.80%), with interproximal surfaces presenting most carious lesions (94.2%). When compared to previous photographs in the literature, caries in anterior teeth appear similar to those described here (e.g., Figure 15 in Schultz, 1935; Figure 5 in Smith et al., 1977; Figure 4 in Stoner, 1995; Figure 5 in Lovell, 1990). Therefore, in addition to the species studied here, other frugivorous primates (including platyrrhine and catarrhine species) seem to display ICAT lesions (see Coyler, 1936; Lovell, 1990; Schultz, 1935; 1956; Smith et al., 1977; Stoner, 1995).

**Figure 4.**
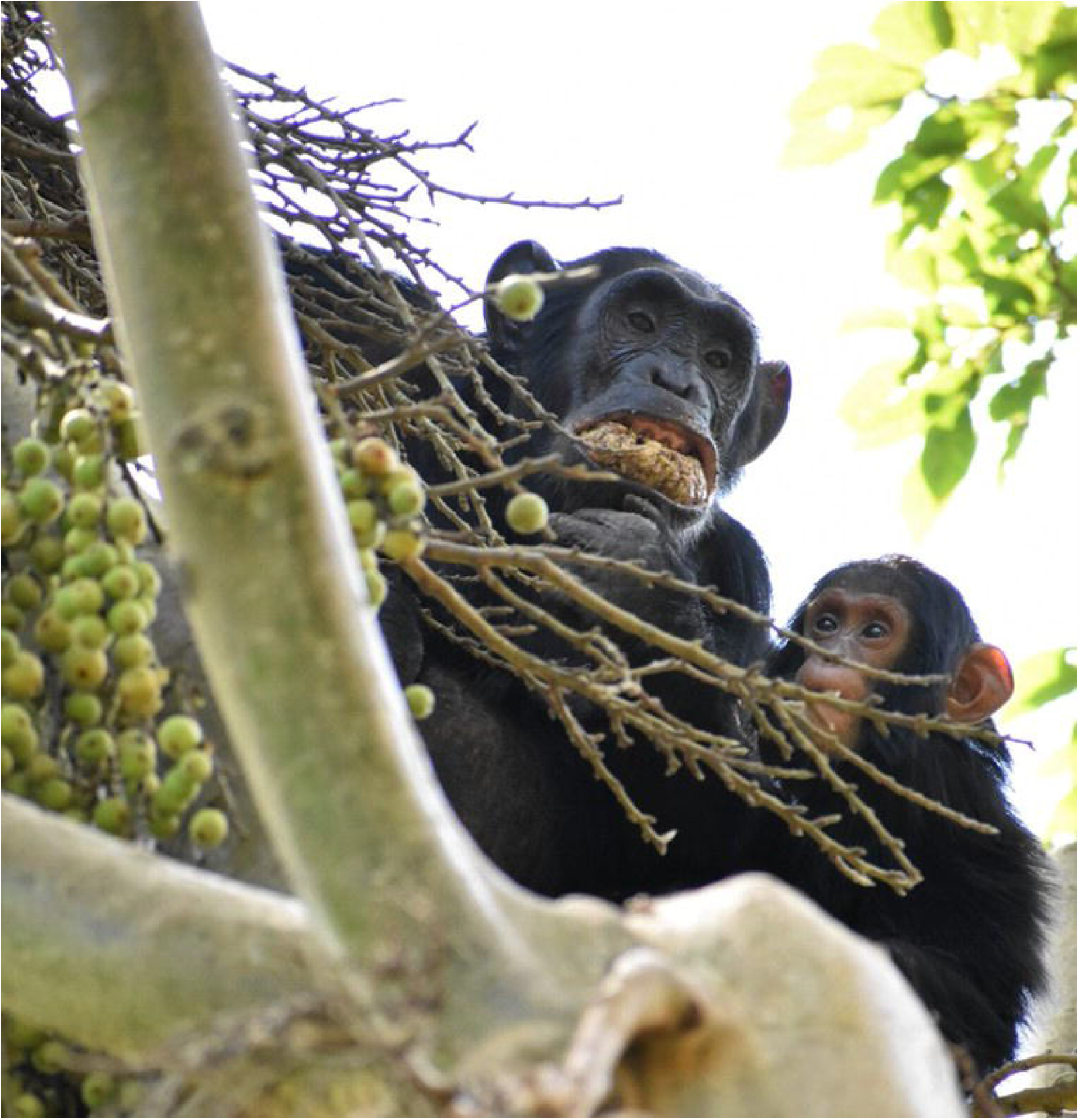
An adult female chimpanzee in Kibale National Park, Uganda, eating figs (*Ficus sur/capensis*) by creating a wadge in the anterior dentition.

In contrast, ICATs seem rare in captive primates, with posterior teeth commonly displaying carious lesions (Anderson & Arnim, 1937; Bowen, 1968; Cohen & Bowen, 1966; Colyer, 1936). Such primates are often fed a cariogenic diet, but have higher caries rates in the posterior teeth (Anderson and Arnim, 1937; Bowen, 1968; Cohen and Bowen, 1966). For example, Colyer (1936) found that almost 90% of carious teeth in captive monkeys were molars. These observations suggest the way in which foods are processed can contribute to the generation of incisor lesions. In the present study, most ICATs were associated with significant attrition/abrasion on the occlusal and interproximal surfaces. Tooth wear, along with periodontal disease and continuous eruption of teeth, may create excessive space for food and bacteria to accumulate below the crown between incisors. Additionally, heavy wear can be associated with root surface exposure, through continuous tooth eruption or periodontal disease, facilitating root caries formation (Hillson, 2008). Many sugary fruits are also acidic, which might create a microenvironment that facilitates proliferation of cariogenic bacteria.

Support for this multifactorial hypothesis in which multiple factors lead to ICAT formation, is found in behavioral observations of wild chimpanzees. Chimpanzees tend to use their lips in tandem with large broad spatulate incisors to process fruits and plants (Hylander, 1975; Lambert, 1999; McGrew, 1999; Suzuki, 1969; Ungar, 1994; van Casteren et al., 2018). Figs have a high concentration of sugars and other carbohydrates and require substantial mechanical processing, which usually involves anterior teeth (Lambert, 1999; Wrangham et al., 1993). Figs are consumed by most chimpanzees (e.g., Basabose, 2002; Matthews et al., 2019; Nishida and Uehara, 1983; Potts et al., 2011; Tweheyo, and Lye, 2003; Watts et al., 2012), often involving holding the chewed fruits in the front part of the mouth, in a behavior called ‘wadging’ (Lambert, 1999; Nishida et al., 1999). Chimpanzees then suck the sugary liquids from the wadge, much of which will sieve through interproximal surfaces of anterior teeth (Figure 4), likely creating a cariogenic microenvironment. Chimpanzees often wadge figs with higher concentrations of sugars (i.e., Ficus sur/capensis, Ficus mucuso) (Danish et al., 2006; Wrangham et al., 1993). Other cariogenic items are also wadged, including honeycombs (Nishida et al., 1999).

Although gorillas had lower ICAT rates than chimpanzees, they still showed these characteristic lesions. Gorillas also regularly eat fruits, many of which are high in soluble sugars (Remis et al., 2001). The cercopithecidae species with ICATs (Dent’s mona monkeys, blue monkeys, and raffles’ banded langurs) are all frugivores that process foods high in fermentable carbohydrates with their anterior dentition. However, the specific types of foods that contribute to lesions formation likely vary. Carious lesions in raffles’ banded langurs may relate to a diet of seeds high in carbohydrates (i.e., starch), processed using incisors (Davies and Bennett, 1988). This process could have led to not only high caries rates in the anterior dentition, but also to high rates of tooth chipping (Towle and Loch, 2021). Dent’s mona and blue monkeys also eat substantial quantities of different fruits (Olaleru, 2017; Takahashi et al., 2019). Further research is needed to ascertain which foods may contribute to ICATs in these species. For example, blue monkeys show substantial variation in feeding ecology across their range (Tesfaye et al., 2013), meaning that a study of caries in populations with detailed dietary and behavioral record is important to elucidate the processes leading to ICAT formation.

Sex differences in caries prevalence are also important, since differences have been observed in humans and other great apes (Lanfranco and Eggers, 2012; Lukacs and Largaespada, 2006; Lukacs, 2011; Stoner, 1995). Dietary and behavioral differences are known between male and female chimpanzees (Gilby et al., 2017; Nakamura et al., 2015; Wrangham and Smuts, 1980). Pregnancy and differences in oral pH, saliva, physiology, life history, and microbiome between the sexes may also be predisposing factors (e.g., Fuss et al., 2018; Lukacs and Largaespada, 2006; Stoner, 1995). The results of the present study support other research suggesting sex differences in caries rates among the great apes, although the present sample of gorillas did not show significant differences.

Recent literature has shown caries is not as rare as previously thought in fossil hominin and extant great apes (e.g., Arnaud et al., 2016; Lacy, 2014; Lacy et al., 2012; Lanfranco & Eggers, 2012; Liu et al., 2015; Margvelashvili et al., 2016; Miles & Grigson, 2003; Stoner, 1995; Towle et al., 2019; Trinkaus et al., 2000). There is also growing evidence that caries was common in other extinct primates (e.g., Fuss et al., 2018; Han and Zhao, 2002; Selig and Silcox, 2020). This study adds further evidence that caries has likely been relatively common in many primate lineages. Additionally, lesion position in the dentition may offer insights into diet and behavior of extinct species, based on comparisons with extant primates. In particular, because of ICATs’ uniform appearance in multiple frugivore species, these may be useful for behavioral interpretations. Given the difference in female and male chimpanzees, caries prevalence may also shed light on sexually dimorphic feeding behaviors and physiology in past primate populations.

## Acknowledgments

We thank I. Livne from the Powell-Cotton Museum and the Study Material Committee from the Primate Research Institute (PRI), Kyoto University, for access to their collections, and T. Ito for assistance during data collection. Research was supported by a Sir Thomas Kay Sidey Postdoctoral fellowship from the University of Otago to Ian Towle and a studentship from Liverpool John Moores University, and was partly performed under the Cooperative Research Program of the PRI (2019-C-20).

## Figure legends

**Supplementary Table 1.** Geographic origin of the samples studied, along with information on the number of individuals split by sex.

| Species | Common name | Location | Total individuals | Males | Females | Unspecified |
| --- | --- | --- | --- | --- | --- | --- |
| *Pan troglodytes* | Chimpanzee | Cameroon (between 3.75-4.25 degrees North; 13.75-14.25 degrees East) | 109 | 41 | 65 | 3 |
| *Gorilla gorilla gorilla* | Western lowland gorilla | Cameroon (between 3.75-4.25 degrees North; 13.75-14.25 degrees East) | 83 | 35 | 46 | 2 |
| *Hylobates klossii* | Kloss's gibbon | Siberut Island, Sumatra, Indonesia | 15 | 0 | 0 | 15 |
| *Simias concolor* | Pig-tailed langur | Mentawai islands (Malibakbak; Silabu; Samekmek; Pasakiat), West Sumatra, Indonesia | 20 | 9 | 11 | 0 |
| *Papio hamadryas* | Hamadryas baboon | Mojo, Ethiopia | 20 | 2 | 7 | 11 |
| *Macaca fuscata* | Japanese macaque | Japan (Chiba; Koshima Island; Yakushima Island) | 48 | 18 | 22 | 8 |
| *Cercopithecus denti* | Dent's mona monkey | Democratic Republic of the Congo (Kalehe; Waikale; Kimbala; Kakolokelwa) | 10 | 5 | 5 | 0 |
| *Cercopithecus mitis* | Blue monkey | Walikale, North Kivu Province, Democratic Republic of the Congo | 8 | 3 | 5 | 0 |
| *Mandrillus spp.* | Mandrill | Cameroon | 10 | 6 | 2 | 2 |
| *Presbytis femoralis* | Raffles' banded langur | Lubuk Jantan (near Koto Alam), West Sumatra, Indonesia | 8 | 3 | 5 | 0 |
| *Presbytis potenziani* | Mentawai langur | Simaileppet Island, West Sumatra, Indonesia | 8 | 5 | 3 | 0 |

**Supplementary Table 2.** Number of individuals with and without caries, split by sex.

| Species | Female |  |  |  | Male |  |  |  |
| --- | --- | --- | --- | --- | --- | --- | --- | --- |
|  | Caries | No caries | Total individuals | % with caries | Caries | No caries | Total individuals | % with caries |
| Blue monkey | 3 | 2 | 5 | 60.00 | 3 | 0 | 3 | 100.00 |
| Dent's mona monkey | 3 | 2 | 5 | 60.00 | 4 | 1 | 5 | 80.00 |
| Mandrill | 0 | 5 | 5 | 0.00 | 1 | 5 | 6 | 16.67 |
| Raffles' banded langur | 3 | 2 | 5 | 60.00 | 1 | 2 | 3 | 33.33 |
| Mentawai langur | 0 | 3 | 3 | 0.00 | 0 | 5 | 5 | 0.00 |
| Japanese macaque | 1 | 21 | 22 | 4.55 | 4 | 14 | 18 | 22.22 |
| Pig-tailed langur | 0 | 11 | 11 | 0.00 | 0 | 9 | 9 | 0.00 |
| Hamadryas baboon | 3 | 15 | 18 | 16.67 | 0 | 2 | 2 | 0.00 |
| Chimpanzee | 26 | 39 | 65 | 40.00 | 7 | 34 | 41 | 17.07 |
| Western lowland gorilla | 3 | 43 | 46 | 6.52 | 4 | 31 | 35 | 11.43 |

## Notes

### Competing Interest Statement

The authors have declared no competing interest.

